# p130Cas contributes to cellular mechanosensing and force exertion

**DOI:** 10.1101/371021

**Authors:** Hedde van Hoorn, Dominique M. Donato, H. Emrah Balcioglu, Erik H. Danen, Thomas Schmidt

## Abstract

Cell survival, differentiation, and migration are all dependent on the cell’s interaction with its external environment. In addition to chemical cues, cells react to their physical environment, particularly the stiffness of the substrate. In order for cells to react to these elements, they must make use of cellular machinery to signal changes in their microenvironment. One such proposed machinery is the protein p130Cas, which has been shown to regulate focal adhesion turnover, actin dynamics, and cell migration. Here we show that p130Cas localizes to focal adhesions depending on substrate stiffness and subsequently modulates cellular force exertion. We compared on substrates of tunable stiffness knock-out CAS^-/-^cells to cells re-expressing either the full-length p130Cas or a mutant lacking the focal adhesion targeting domains. On polyacrylamide gels, we observed that p130Cas prevented focal adhesion formation at low stiffness. On structured micro-pillar arrays, p130Cas preferentially localized to sites of force exertion when the apparent Young’s modulus of the substrate was higher than E = 47 kPa. Stiffness-dependent localization of p130Cas coincided with slower, but increased force exertion for the full-length p130Cas. Cas localization to focal adhesions preceded force build-up by three minutes, suggesting a coordinating role for p130Cas in the cellular mechanoresponse. Thus, p130Cas appears to relay mechanosensory information in the cell through its ability to tune force exertion at the focal adhesion.

## Introduction

Tissues, cells and extracellular matrices vary greatly in stiffness (1; 2). In recent years, it has become apparent that this stiffness not only emerges from, but also dictates biological function (3; 4; 5; 6). Stiffness-dependent cellular behaviour is attributed to a variety of molecular responses, such as transcription factor incorporation (7), remodelling of the nuclear envelope (3), and differential focal adhesion (FA) protein signalling (8). FAs are of particular interest in this respect, as they are the main sites of cellular force transmission on the extracellular matrix. FAs are multi-molecular complexes that contain a multitude of functional biological interactions (9). The combination of force transmission and biological functionality observed at these sites has given rise to a variety of hypotheses on the stiffness-dependent mechano-sensory role of FA proteins. On substrates of compliant surfaces, cell spread and FA size have been shown to increase in a mechano-sensitive way and have furthermore been linked to various signalling molecules in FAs (10; 11). Here, we address the mechano-sensory role of one such FA signalling protein, p130Cas.

p130Cas is a member of the Cas (Crk associated substrate) family that regulates cellular behaviours such as migration, apoptosis, cell-cycle progression and differentiation (12). Knocking out p130Cas constitutively leads to embryonic lethality 11.5-12.5 days post-fertilization. Moreover, p130Cas (also known as breast cancer anti-oestrogen resistance protein 1, BCAR1) has been associated with resistance to anti-oestrogen treatment in breast cancer patients (13). Another study revealed that p130Cas over-expression leads to a poor prognosis in non-small-cell lung cancer (14) and is generally of importance in the progression of several cancer types (15).

As a scaffolding protein, p130Cas initiates a multitude of signalling cascades. It is organized into a Src homology 3 (SH3) domain on the N-terminus, a proline-rich domain, a central substrate domain (SD) containing 15 YxxP motifs, a serine-rich domain, a Src-binding domain (SBD) containing motifs for binding both the Src SH3 and SH2 domains, and a well conserved Cas-family C-terminal Homology (CCH) domain. A simplified schematic drawing of the protein structure is shown in Fig.1. p130Cas promotes cell spreading in response to integrin engagement and regulates cell migration through increased FA assembly and turnover (16; 17). Both the SH3 domain on the N-terminus and the CCH domain on the C-terminus localize the protein to FAs (18). Tyrosine phosphorylation of the SD directly influences cell migration, actin dynamics and FA dynamics (19; 18; 16).

**Figure 1:**
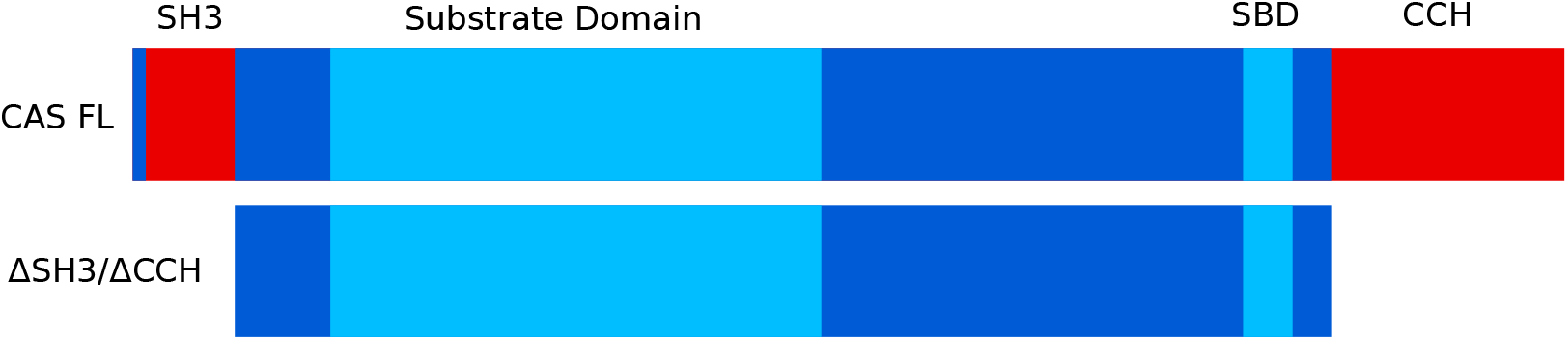
P130Cas structure and domains. The ability of p130Cas to localize to FAs is mediated by the SH3 and CCH domains. The Substrate Domain (SD) contains 15 YxxP motifs that can be phosphorylated by Src. Src kinase can bind through the SBD and FAK via the SH3 domain (18). Both the full-length and ∆SH3/∆CCH variants of p130Cas were used for this study.

In addition to its importance in cellular function and disease, a mechanosensory role has been proposed for p130Cas, which has remained unclear. Experiments on Triton-X extracted cytoskeletons showed an increase in tyrosine phosphorylation of p130Cas when cells were physically stretched (20). Similar data have been obtained *in vitro* with a recombinant p130Cas SD, demonstrating that its intrinsically disordered SD can be unfolded at pN forces (21; 22). This force range is expected to occur at sites of integrins in FAs (23). From the observations it was proposed that a large enough force to physically stretch the protein could increase the likelihood of SD tyrosine phosphorylation. It is unclear, however, whether physiological forces and extracellular stiffnesses have an impact on p130Cas localization to and force exertion at FAs.

Here, we show that p130Cas functions as a mechanosensor in cells within physiological stiffness ranges of relevant tissue, and that it mediates a mechanical-biological-mechanical coupling between the cell and the extracellular matrix. p130Cas only localized to FA sites when the substrate has a Young’s modulus higher than 47 kPa (shear modulus >15 kPa). Subsequently to localization, p130Cas changed cellular force exertion dynamics. On p130Cas localization which only occurred on substrates of elevated stiffness, cells exerted both larger and more persistent forces. Dynamic observations relating p130CAS localization to cellular force generation suggests that p130Cas serves as a physico-chemical coupler directing mechanosensing and mechanotransduction in cells on stiff matrices.

## Materials and methods

### Cell biology

All experiments were performed in cells deficient of endogeneous p130Cas. CAS^-/-^mouse embryonic fibroblasts (MEFs) were used and adapted as previously described (18). Further we used retroviral transduction to obtain CAS^-/-^cells into which the full length p130Cas fused to YFP (CAS^FL^), or a truncated version fused to YFP lacking both the SH3 and CCH domains (CAS^∆SH3/∆SSH^) were stably reintroduced (for details see (18)).

MEFs were grown in high glucose Dulbecco’s modified Eagle medium without phenol red, supplemented with 10% fetal bovine serum, 1% penicillin/streptomycin and 1% glutamax. Cells were maintained in an incubator at 37°C, with 7% CO_2_. On PA gels, cells were seeded 1 hour prior to fixation with 4% paraformaldehyde (PFA) in phosphate-buffered saline (PBS). On micropillar arrays, cells were seeded 6 hours prior to overnight imaging in full medium (since cells were not sufficiently adherent after 1 hour to withstand the pillar inversion process). In both cases, cells were seeded at single-cell density.

### PA gels

PA gels were prepared similarly as previously described (24; 25). Briefly, 12 mm sterile coverslips were placed in 24-well plates, cleaned with 0.1 M NaOH, and then rendered hydrophilic by incubating with 0.5% 3-aminopropyltrimethoxysilane (3-APTMO, Sigma-Aldrich). Coverslips were subsequently washed thoroughly with sterile milliQ water and incubated in 0.5% glutaraldehyde and dried overnight in a laminar flow cabinet. Coverslips of 10 mm diameter were rendered hydrophobic by incubating with a solution of 10% Surfa-Sil in chloroform (Thermo Scientific), washed in 100% chloroform, then in methanol and dried under laminar flow. PA solutions were made with a mixture of acrylamide and bis-acrylamide as given in the supplemental table (Tab.S1).

After mixing acrylamide and bis-acrylamide, 1.5*μ*l TEMED and 5*μ*l of 10% ammonium persulfate were added to start polymerisation in a total volume of 1 ml. On each 12 mm coverslip, 10*μ*l was applied and 10 mm coverslips were placed on the top to make a flat layer and left to polymerise for 1 hour. After 15 minutes washing with 50 mM 4-(2-hydroxyethyl)-1-piperazineethanesulfonic acid (HEPES), the top 10 mm coverslips were removed and gels were washed once with 50 mM HEPES. Cross-links for fibronectin on PA gels were created by incubating gels with 0.5 mM sulfo-syccinimidyl-6-[4’azido-2’nitrophenylamino]hexanoate (sulfo-SANPAH, Thermo Scientific) in 50 mM HEPES under UV light for 8 min. The gels were washed with 50 mM HEPES, then incubated with sulfo-SANPAH again under UV light, and washed extensively with 50 mM HEPES before incubating overnight at 4°C in 10*μ*g/ml fibronectin and 50*μ*g/ml Alexa405-conjugated fibronectin in PBS. After removing the fibronectin solution by washing with PBS, PA gels were allowed to equilibrate for one hour in complete culture media at 37°C before seeding with 2.5 10^4^ cells/well in complete media. Cells were allowed to adhere and spread before fixation and imaging by incubating for one hour at 37°C and 7% CO_2_.

The PA gel stiffness was measured by applying polacrylamide solutions directly to the plate of a rheometer (Anton-Paar MCR 501) during polymerisation, while inducing shear with a 40 mm diameter cone plate geometry to obtain the shear modulus G. Both storage and loss moduli were quantified, confirming a negligible viscous behaviour. Finally, assuming a Poisson ratio of 0.5 for an incompressible material, the Young’s modulus *E* = 3 × *G* of the gel was calculated for proper comparison with the micropillar experiments. The moduli obtained coincide with those reported earlier (24). It should be noted that the stiffness values reported here consistently refer to Young’s modulus, whereas in literature the use of Young’s and shear moduli is scattered.

### Immunostaining

After 1 hour spreading on PA gels, the cells were fixed using 4% PFA in PBS for 15 minutes, permeabilized using 0.1% Triton-X for 10 minutes and blocked using 5% normal goat serum for one hour (all diluted in PBS). Antibodies were diluted in blocking buffer and paxillin was recognized using a mouse-anti-paxillin IgG (BD Biosciences), further recognized by a Cy3-conjugated AffiniPure goat anti-mouse IgG F(ab’)2 antibody (Jackson Immunoresearch). Cells on pillars were fixed using the same method, but stained with mouse-anti-paxillin primary antibody (BD-Transduction Laboratories) and then recognized by an Alexa405-goat-anti-mouse secondary antibody (Jackson Immunoresearch).

### Microscopy

Both live and fixed cell imaging were performed on an adapted Zeiss microscope (AxioVert 200) with a spinning disk confocal unit (Yokogawa X-1) and a back-illuminatedEM-CCD camera (Andor DU-897). Multiple laser lines were combined and controlled using an acousto-optical tunable filter (AOTF, AA Optoelectronics), then coupled into the spinning disk unit using a polarization-maintaining fibre. From the back-port, a home-built focus-hold system was incorporated, using a 850 nm laser diode, dichroic mirror (Chroma) and a photo-diode detector (Thorlabs). A Märzhauser XY-stage controlled automated movement to various positions on the sample at specific focus positions. Imaging was controlled using Andor IQ-software, while the focus-hold and stage movement were controlled using Labview software (National Instruments). The samples were mounted in a specially-designed, stable coverslip holder that fitted directly into the microscope incubator (Tokai Hit). Overnight live-cell imaging was performed at 37°C and 5% CO_2_ with. 12 positions were used per experiment.

### FA analysis

Automated image analysis in Matlab (Mathworks) was developed to quantify FAs and their spatial properties. Using frequency-filtering and edge-detection algorithms, the cell contour and FAs were detected. A threshold relative to background noise was set to detect the cell boundary, which was then set as a binary object. Within this object, the distance to the cell edge was calculated to include only FAs at the periphery. FA objects were detected by a separate thresholding step. We have previously described this algorithm in application to multiple cell types (25). An example of a cell edge and the detected FAs is given in the supplementary figure (Fig.S1).

### PDMS Micropillars

Poly(dimethyl-)siloxane (PDMS, Sylgard 184, Dow Corning) was mixed with 1:10 crosslinker:prepolymer ratio and poured over the silicon master. After 20 hours curing at 110°C, the arrays were peeled off and micro-contact printed with fibronectin and fibronectin labeled with Alexa647 (Invitrogen). Prior to micro-contact printing, the PDMS micropillar array was activated with a UV-Ozone cleaner (Jelight) for 10 minutes. Subsequently, the non-printed parts were passivated with 0.2% pluronic (F-127, Sigma Aldrich) in PBS. The substrates were thoroughly rinsed with PBS, submerged in full media. 10^5^ cells per array were seeded. After 6 hours spreading, the samples were mounted on the microscope in an inverted configuration (as previously described (26)) and imaged overnight. The bending stiffness of the micropillars was previously determined (26) to be: 16.2, 65.8, and 199 nN/um, respectively. The effective stiffness was calculated using a relation derived by Ghibaudo et al. (27; 28). The effective Young’s modulus of E = 12, 47 and 137 kPa was determined for 3.1, 4.2 and 6.9*μ*m high pillars, respectively. A sketch of the experimental realization is given in supplementary.

### Analysis of forces

Cellular forces were quantified by analysis of the pillar deflections as previously described (26; 25). Increase and decrease of transient force dynamics were fitted using a logistic function that describes the force build-up at a given rate up to a maximum size (eq. 1). In general, the dynamics will be characterised by two independent rates for build-up and break-down. As those appeared indistinguishable in our experiments (i.e. there was no significant difference between the build-up and break-down rate), we assumed that both rates are given by a single rate, *r*. The resulting functional form for *F*(*t*), as given by eq. 2, (including a time offset) was fitted to the dynamic force data to yield *F_max_* and *r*. Finally, we determined the time point, *t*_1/2_, at which half the maximal force was reached (eq. 3).

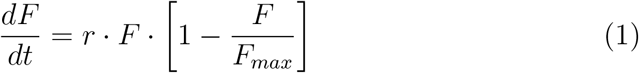

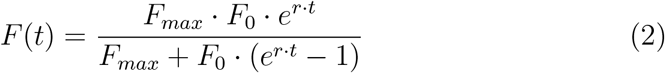

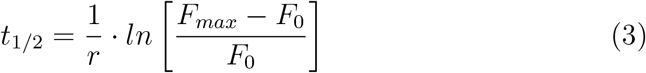

Half-times of transient force-increase on a single pillar and simultaneous fluorescence increase of p130Cas were quantified using specifically designed Matlab (Mathworks) scripts. The time difference between force exertion and p130Cas localization was denoted as ∆*t*.

### Statistics

Statistical analysis on all data was performed using two-sample Kolmogorov-Smirnov tests. p-values for significance in difference of the values were calculated. In figures p *<* 0.05 is depicted by (*), p *<* 0.01 by (**), and p *<*0.001 by (***), respectively. Details on e.g. number of cells are found in supplementary tabels (Tab.S2&S3).

## Results

### Stiffness-dependent focal adhesion formation is regulated by p130Cas

We hypothesized that the previously observed p130Cas-mediated cell spreading and focal adhesion (FA) turnover were stiffness-dependent (29; 16; 17). Therefore, we compared p130Cas-knockout (CAS^-/-^) mouse-embryonic fibroblasts (MEFs) with CAS^-/-^cells into which the full-length p130Cas has been re-expressed (CAS^FL^) on their initial spreading behaviour on polyacrylamide (PA) gels. The PA-gel stiffness was varied by changing the ratio of acrylamide:bis-acrylamide, yielding a Young’s modulus that ranged from 28 to 222 kPa (shear modulus range of 9.3 to 74 kPa). The range was chosen since it covers the range of typical tissues stiffnesses. After 1 hour of spreading, cells were fixed and stained for the FA protein paxillin. Cell and FA area were quantified using specifically designed algorithms (for details see Materials).

On the stiffest PA gels, both cell lines displayed a morphology similar to what has been reported on glass (29) (see Fig.2A-B). Both CAS^FL^and CAS^-/-^cells spread poorly on the softest substrates (Fig.2). A significant increase in cell spreading area was observed between 42-87 kPa for CAS^-/-^, and between 87-222 kPa for CAS^FL^(Fig.2C). With the increase in spreading area, an increase in FA-area was observed for both cell lines. The FA-area increased between 42-87 kPa for CAS^-/-^(Fig.2D), in parallel with the increase in spreading area. In contrast CAS^FL^did not develop detectable FAs on the lowest stiffness substrates (28 kPa), which only became visible on gels of 42 kPa. Beyond this threshold FA-area remained constant. Altogether, our findings indicate that expression of p130Cas shifts the onset of cell spreading to higher stiffness, and restricts the formation of FAs on substrates of low stiffness.

**Figure 2:**
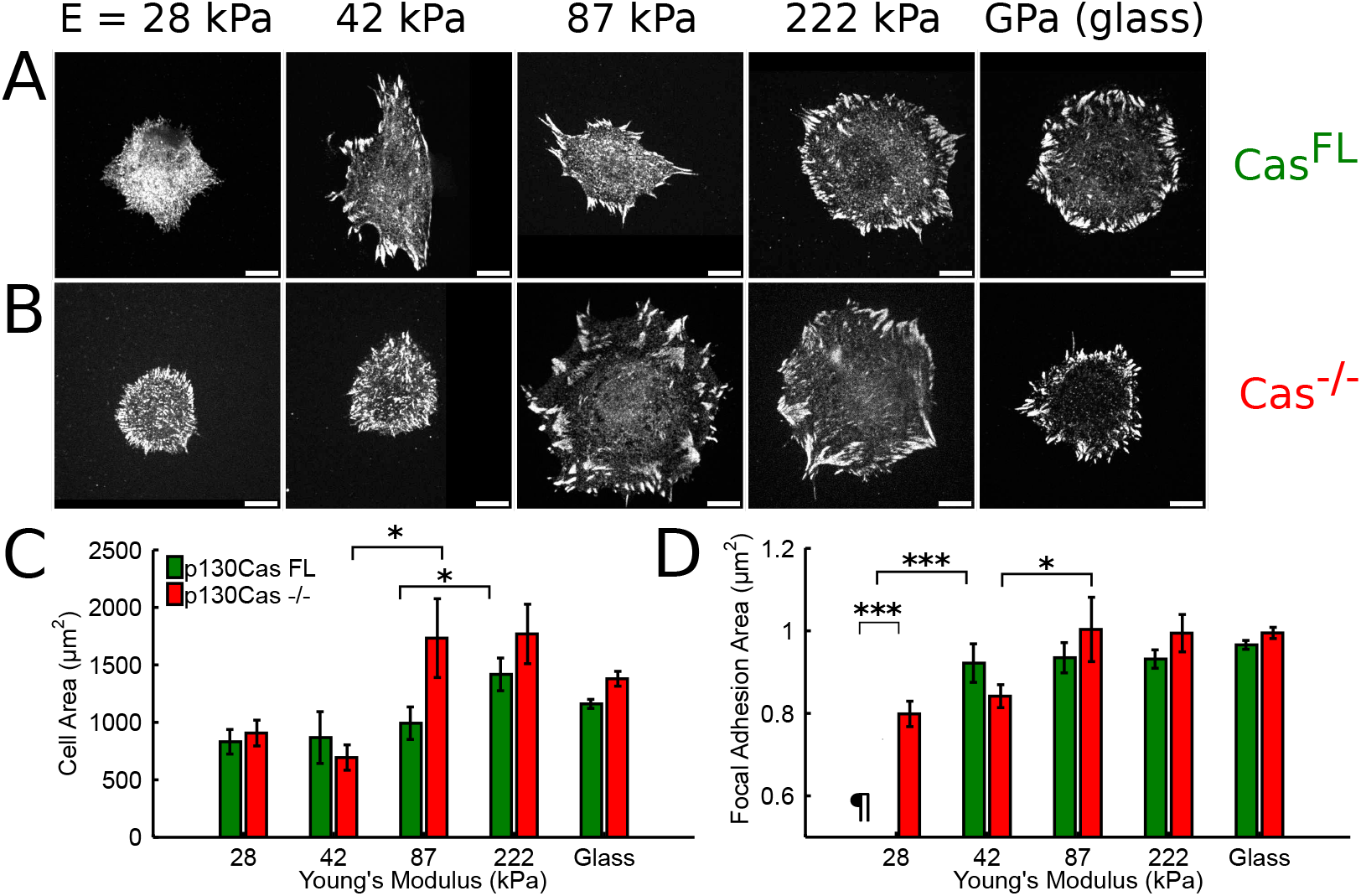
P130Cas expression influences stiffness-dependent cell spreading and focal adhesion formation. (A) CAS^FL^, and (B) CAS^-/-^mouse embryonic fibroblasts were seeded on polyacrylamide gels of varying stiffness (Young’s modulus indicated in kPa) and on glass. All surfaces were coated with fibronectin for 1 hour prior to fixation. FAs were labelled by immunostaining for paxillin. Means and s.e.m for (C) cell-area, and (D) FA-area were quantified for both cell types. No FAs were found for CAS^FL^cells on 28 kPa substrates (denoted by *¶*). Scale bars in (A) and (B) are 10*μm*. Data were analyzed on ≥ 750 FAs from ≥ 11 cells per stiffness, and for ≥ 2 independent experiments.

### P130Cas localization to FAs depends on stiffness

Since p130Cas influenced stiffness-dependent FA formation, we hypothesized that p130Cas may localize to sites of force exertion in a stiffness-depending manner. To test our hypothesis we performed experiments on deformable micropillar arrays. Micropillar arrays allowed us to measure the local cellular forces, while controlling the global, cell-wide extracellular stiffness. The sub-cellular fluorescence localization of p130Cas-YFP was quantified simultaneously with the force measurements on micropillars.

We tuned the global stiffness of the micropillar arrays by varying the height of the micropillars while maintaining a pillar diameter of 1.9-2.1 *μ*m with interpillar spacing of 2 *μ*m. Micropillar arrays with a height of 6.9 *μ*m, 4.1 *μ*m and 3.2 *μ*m, resulted in effective Young’s moduli of 12, 47 and 137 kPa, respectively (Fig.S2). Pillar deflection from the expected hexagonal pillar-grid pattern was determined to an accuracy of 30 nm (26), which translated to a force resolution of 0.24, 0.98, and 3.0 nN for the 6.9, 4.1, and 3.2 *μ*m arrays, respectively.

Cells attached only to the tops of micropillars and spread fully 6 hours after seeding. FA formation occurred on pillar-tops, as we and others previously observed (28; 31; 26). Significant forces were measured within FAs only (Fig.S3 for paxillin immunostaining). Forces exerted on micropillars were in the range of 1 - 60 nN. We observed the localization of p130Cas simultaneously with local cellular force exertion. Fig 3A shows pillars in red, p130Cas fluorescence in green, micropillar deflections as white arrows and p130Cas fluorescence in separate grayscale images.

**Figure 3:**
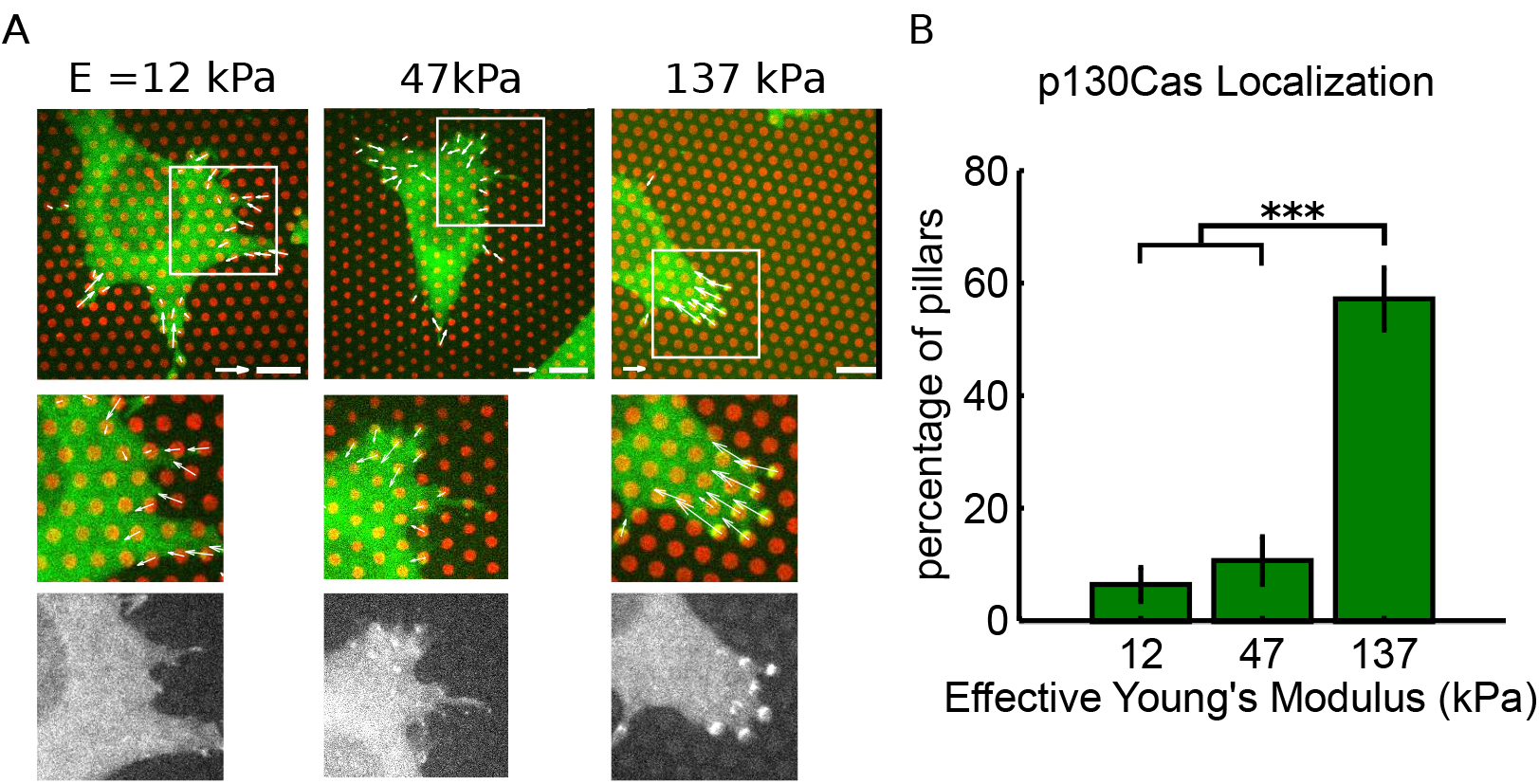
p130Cas localization to sites of force exertion is dependent on substrate stiffness. (A) CAS^FL^expressing p130Cas-Venus (green) at endogenous levels on micropillars (red) of varying stiffness. Insets (midle and bottom) are magnified by 1.75. Forces higher than the detection limit are displayed as white arrows. Grayscale images (bottom) show p130Cas fluorescence only. Scale bars corresponds to 10 *μ*m. Force scale bar corresponds to 20 nN (12 kPa and 47kPa), and 50 nN (137 kPa), respectively. (B) Percentage of deflected micropillars per cell to which p130Cas fluorescence was co-localized (mean ± s.e.m.). Data were obtained from ≥18 cells with ≥260 pillars measured per stiffness, from a total of ≥2 experiments.

p130Cas localized to sites of force exertion predominantly on stiffer micropillar arrays. The fraction of pillars that showed p130Cas localization for the three stiffnesses is given in Fig.3B. On 12 and 47 kPa micropillar arrays, 5-10 % of the deflected pillars showed p130Cas localization. At a global stiffness of 137 kPa, a significant increase in p130Cas localization was observed. At ≈60% of the attachment sites where more than 30 nN of force was applied, p130Cas was localized to pillars. p130Cas localization on 137 kPa micropillars closely resembled its localization to FAs on glass substrates (18).

These results demonstrate that p130Cas localized to force exertion sites depending on stiffness. The increased localization of p130Cas in FAs on the pillar arrays was observed in a similar stiffness range as the initiation of FA area increase on PA gels (fig.2). Hence we conclude that p130Cas responds to variations in the global extracellular matrix stiffness rather than to the local stiffness represented by a single pillar.

### Force exertion dynamics depend on substrate stiffness and p130Cas localization

Since p130Cas alters FA dynamics on glass (18; 16), and given the mechanosensitive function of p130Cas in CAS^FL^cells, we hypothesized that p130Cas localization to FAs may further influence force exertion itself. We therefore investigated the dynamics of force exertion on micropillar arrays of varying effective stiffness. Fig.4 shows still images from a time series movie of p130Cas-YFP fluorescence (green) and the tops of micropillars (red). As expected, cells displayed random migration with an extended leading edge (see supplemental movies for cell behavior on micropillars of varying stiffness). Transient contractile forces opposite to the direction of migration were visible at the leading edge of cells.

**Figure 4:**
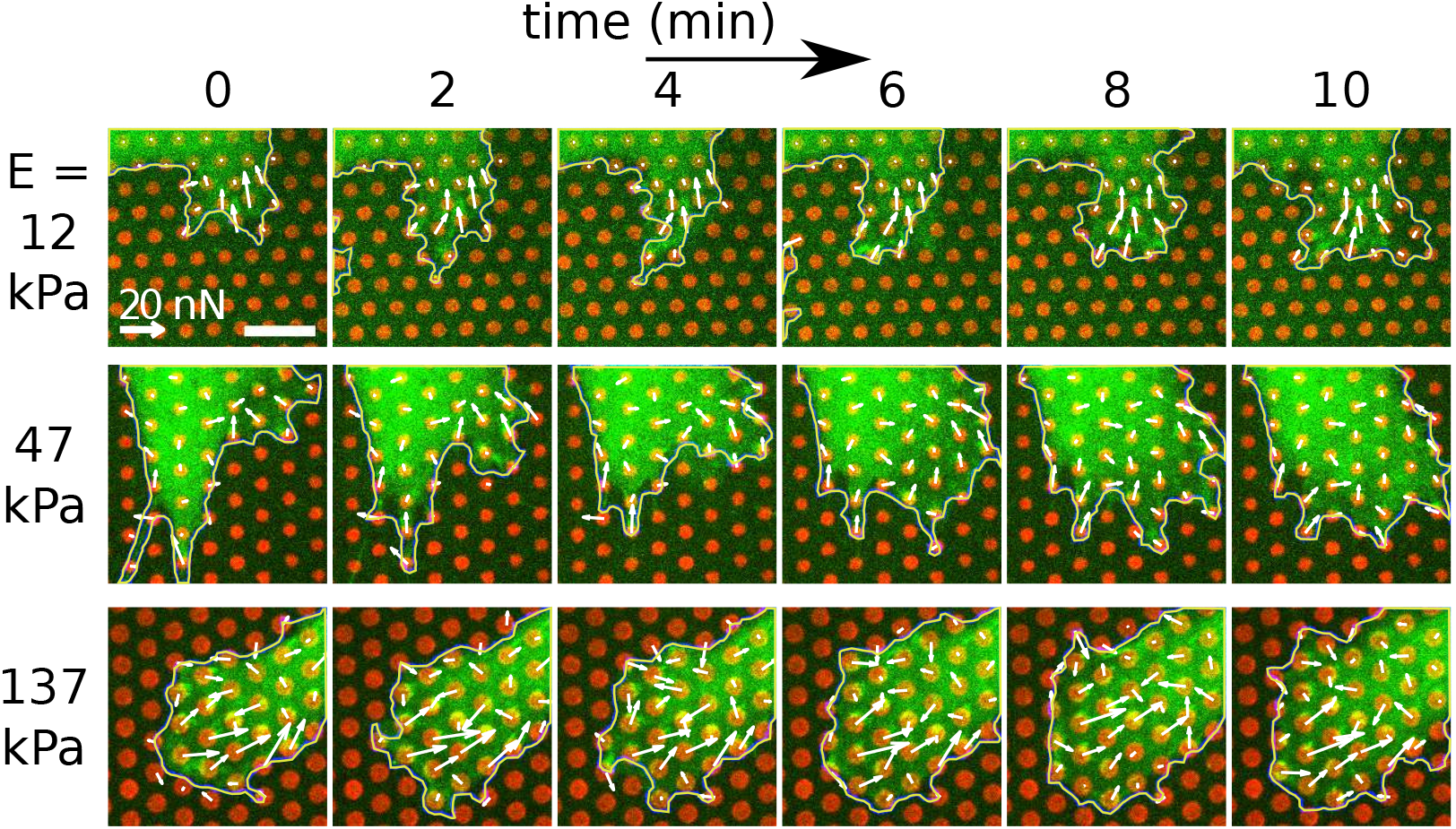
Time-lapse imaging of force exertion and p130Cas localization. CAS^FL^expressing p130Cas-Venus YFP (green) at endogenous levels were imaged every 2 minutes to examine the dynamics of force exertion on micropillars (red). Pillar deflections (white arrows), and the cell edge (yellow) are plotted. (image scale bars 10 *μ*m; force scale bars 20 nN).

To test the effect of p130Cas localization to sites of force exertion, we quantified the force exertion characteristics of CAS^-/-^cells either re-expressing the full p130Cas-YFP protein (CAS^FL^), or a double-deletion mutant lacking both the SH3 and CCH domains (CAS^∆SH3/∆SSH^). The latter mutant was previously characterized and was shown to lack FA localization, to have reduced SD tyrosine phosphorylation, and displays reduced cell migration (18). Here we confirmed by fluorescence localization that CAS^∆SH3/∆SSH^was unable to localize to FAs (Fig.S5). As shown in fig.3 and fig.4 CAS^FL^did localize to FAs on high-stiffness pillars.

Typical curves of the force development over time for individual pillars are shown in Fig.5A (blue dots) for the different micropillar stiffnesses. The transient force dynamics was modelled with a logistic function (see Materials for details) that was characterized by two independent parameters: (i) the maximum force *F_max_*, and (ii) the relative rate 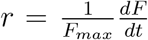 by which *F_max_* is approached. We found that force build-up and force decay rates did not differ significantly, hence the model contained only one characteristic relative force rate. This minimal model provided a good fit to the data, as shown in Fig.5A (red line for the force increase and green line for the force decrease).

**Figure 5:**
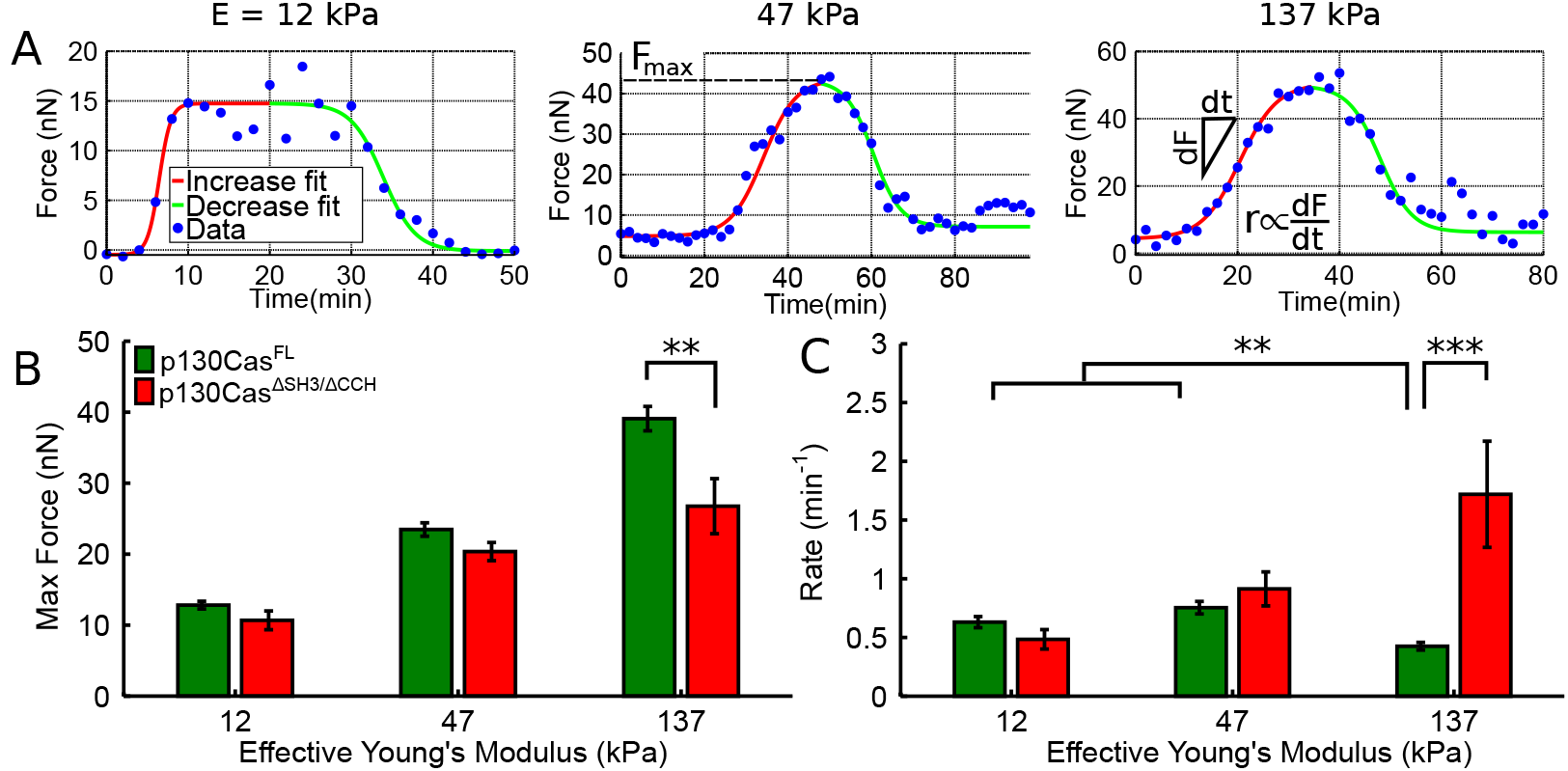
Quantification of p130Cas-dependent force exertion dynamics. Transient force exertion on individual pillars was quantified by fitting increase and decrease curves to a logistic function. (A) Examples for CAS^FL^. From these data the maximal force F_*max*_, and the rate of force exertion. were obtained. (B) *F_max_* as a function of substrate stiffness (green: CAS^FL^; red: CAS^∆SH3/∆SSH^). (C) Dependence of the force rate. for different substrate stiffness. Bars represent mean±s.e.m. obtained from ≥2 experiments with ≥12 cells and ≥110 pillar deflections each.

The maximal force F_*max*_ and force rates *r* are shown in Fig.5B-C for different stiffness of the micropillar arrays and for CAS^FL^(green) and CAS^∆SH3/∆SSH^(red) cells, respectively. In CAS^FL^cells *F_max_* increased with effective substrate from 12.8±0.5 to 39±2 nN when the substrate modulus was increased from 12 to 137 kPa (Fig.5B). A likewise increase was observed for CAS^∆SH3/∆SSH^(Fig.5B), though the overall force values were lower. On 137 kPa micropillars *F_max_* was 27±4 nN for the CAS^∆SH3/∆SSH^cells, ≈10 nN less than what was observed for CAS^FL^cells.

While the rate of force exertion for CAS^FL^cells was independent on substrate stiffness below 47 kPa with 0.75±0.03 min^−1^, it significantly decreased to 0.45±0.03 min^−1^ on 137 kPa micropillars (Fig.5C). This observation was opposite to the behavior for cell and focal adhesion area shown in Fig.2, and the increase in F_*max*_ in Fig.5C. In contrary, the rate of force exertion for CAS^∆SH3/∆SSH^steadily increased from (0.8±0.2) to (1.72 ±0.45) min^−1^ when the substrate stiffness was increased, paralleling the behavior observed for F_*max*_ (Fig.5C). From our observations we conclude that the presence of functional p130Cas in FAs (Fig.3), correlates with an increased force exertion with substrate stiffness as a consequence of an increase in the maximal force and a decrease in force rate. Loss of p130Cas functionality in CAS^∆SH3/∆SSH^led to lower cellular forces as a result in reduced maximal force and a higher force rate.

Since p130Cas localized to FAs more effectively on substrates of higher stiffness, we deduced that localization may directly change local force exertion. Our experiments, as detailed in the previous paragraph, allowed us to further examine the cause and effect of p130Cas localization to FAs by simultaneously quantifying the time-course of p130Cas fluorescence and the magnitude of force on a single micropillar. Since p130Cas fluorescence was only significantly localized to137 kPa pillars, we further investigated the correlated dynamics of p130Cas localization and force exertion on the stiffest substrates in CAS^FL^MEFs.

Fig.6A shows the time-dependent force development (blue) and the integrated p130Cas fluorescence on top of one pillar (red). Quantitative analysis of Cas-YFP fluorescence in a FA on the top of a single pillar is given in the supplemental material (Fig.S4). As indicated in Fig.6A by the time points at which the fluorescence and force signals reached their half-maximal values, p130Cas localization preceded the build-up of cellular force. In the data shown in Fig.6A the time lag between fluorescence signal and force signal was ∆t = 3 min. From the analysis of ∆t for 44 pillars in 3 cells we constructed the probability density of this time delay (Fig.6B). The probability density reached a maximum at (3.2±0.7) min, the most probable time that p130Cas localization preceded force exertion. Hence, our data suggest that p130Cas has a significant role in FA organization which in turn regulates the rate and extent of cellular force exertion.

**Figure 6:**
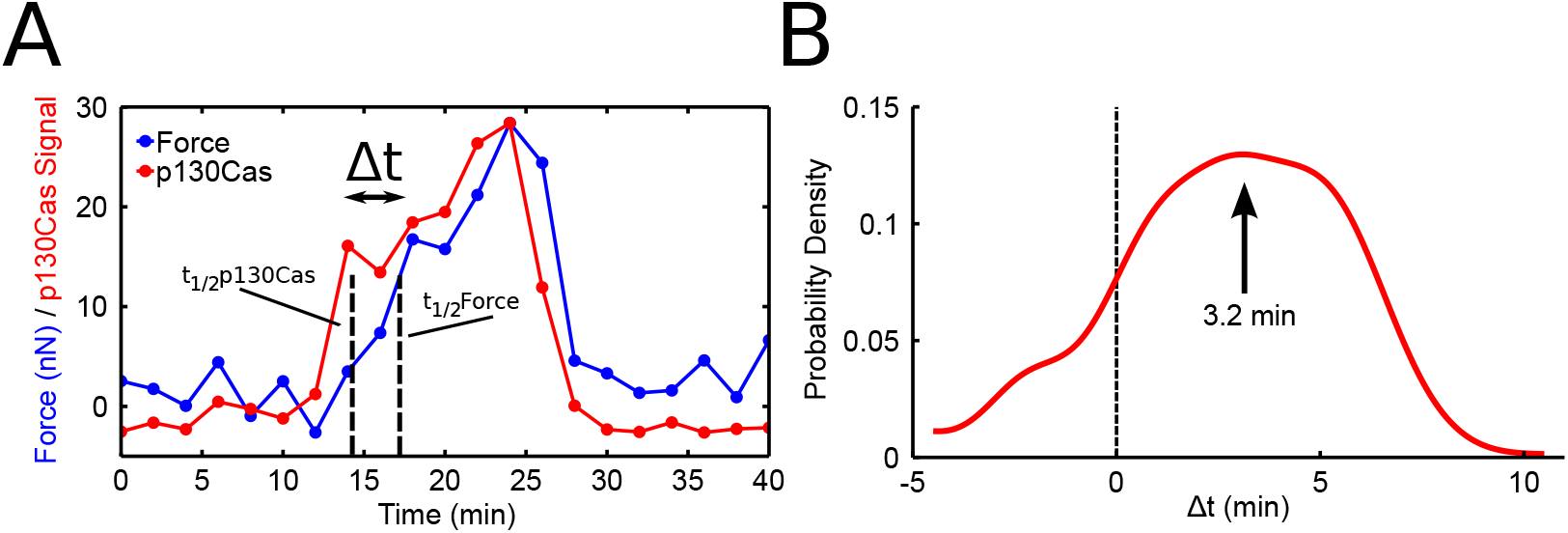
Force exertion and p130Cas localization on an individual pillar. (A) Typically, p130Cas fluorescence was detected prior to force exertion. The halftimes (t_1/2_) are indicated by dashed lines. (B) Probability density of the time lag ∆t between p130Cas localization and force exertion. The p130Cas localization accumulated (3.2±0.7) minutes before force exertion. Data were taken from 44 FAs/curves from a total of 3 cells.

## Discussion

Reconstitution and stretch experiments *in vitro* have highlighted a potential role for p130Cas as a cellular mechanosensor (20; 21; 22). This hypothesis was furthermore supported by *in vivo* experiments that showed that p130Cas altered focal adhesion formation and dynamics (16; 17; 18) on soft substrates. However, a description of how p130Cas may sense extracellular stiffness and in which way protein interactions in cells may influence force exertion is so far unclear.

Here, we showed that p130Cas has a stiffness-dependent role in cellular mechanosensing. p130Cas alters FA formation depending on extracellular stiffness. We found that p130Cas alters FA formation and only localizes to FAs when the extracellular matrix stiffness was larger than a threshold value of 47 kPa. Since this response was observed both on homogeneous PA gels and on pillar arrays which were structured on the micrometer scale, we concluded that p130Cas senses the global extracellular stiffness on the cellular scale. We further found that localization of p130Cas depends on how much force is exerted by the cell. The latter finding makes it conceivable that p130Cas responds to extracellular stiffness by acting as a mechanosensory relay point for cellular force exertion feedback.

Our results, together with the signalling pathways downstream of p130Cas described in the literature, make it likely that p130Cas is one of the players in cellular stiffness responses. After p130Cas localises to FAs, it will interact with various proteins to initiate a functional response. It has recently been noted that tyrosine phosphorylation of p130Cas, as well as paxillin, vinculin, and FAK are upregulated on stiffer substrates as compared to low stiffness substrates (34). Upregulation was attributed to FAK/p130Cas/Rac mediated stiffening of the actin cytoskeleton in a Src-dependent manner. Independently, tyrosine phosphorylation of the p130Cas SD has been suggested as a physical link between the actin cytoskeleton and the FA thereby regulating actomyosin contractility in migrating cells (35). Phosphorylation of p130Cas can thus regulate the amount of force exertion and the rate of force exertion mediated by the Rac pathway. Here we observed that localization of p130Cas to FA sites preceded force exertion by ~3 minutes (see Fig.6). This timescale corresponds to the timescale of typical FA protein phosphorylation events, and to phosphorylation of p130Cas in particular (36). Hence, it appears likely that after FA localization, p130Cas is force-activated, leading to the phosphorylation events which relay the mechanical signal into a biochemical signal.

p130Cas has many binding partners, including FAK and vinculin, which both have significant roles in FA dynamics and in mechanosensing. In terms of mechanosensing, the SH3 domain of p130Cas plays an important role in localization to FAs, which is known to occur through FAK and vinculin binding (18; 37). Previously, it was noted that p130Cas must localize to FAs and interact with vinculin in order to exert maximal traction forces on soft PA gels (37). This interaction may be critical for the initial sensing of the substrate stiffness and the determination as to whether or not p130Cas will localize to FAs. Alternatively, the SD provides a physical link (through its proposed stretching mechanism), to direct signaling cascades that promote force exertion (actomyosin contractility and FA protein phosphorylation/turnover). Enhanced actin polymerisation via p130Cas/Crk coupling is also likely to have an effect on the increase in force generation that was observed with enhanced p130Cas localization on hard pillars. Indeed, p130Cas/Crk coupling has long been noted to enhance actin dynamics, which is also thought to be important in mechanotransduction (38; 16). Interplay between p130Cas and other FA proteins implicated in mechanotransduction (such as vinculin and talin) is likely involved in these processes. Therefore, p130Cas is ideally poised at the intersection of multiple signalling pathways to influence mechanosensing.

Our mechanical data are suggestive of earlier biochemical results that identified the formation of specific FA-protein interactions with p130Cas, leading to increased Rac activity (34). On glass, p130Cas has been shown to strongly influence actin polymerization and FA formation (16; 18). Here, we observed an increase in extracellular force exertion when p130Cas was present in FAs that likewise triggered further p130Cas localization. The increased presence of p130Cas can thus clarify why the total force exerted is not only larger, but also more persistent in comparison to the case where p130Cas was not present or mutated. In this way, p130Cas might have a crucial function in local stabilisation of the FA. In stabilized FAs, p130Cas might further enhance recruitment of signalling partners to promote larger and more persistent forces. A higher persistence in force exertion was observed as a decrease in the force exertion rate. This finding is suggestive of p130Cas acting as a mechanosensor both in outside-in signalling (its localization depends on stiffness), and in inside-out signalling (it influences cellular force exertion). An overview of our proposed sensing (outside-in) and responding (inside-out) mechano-chemical coupling by p130Cas is summarized in Fig.7.

**Figure 7:**
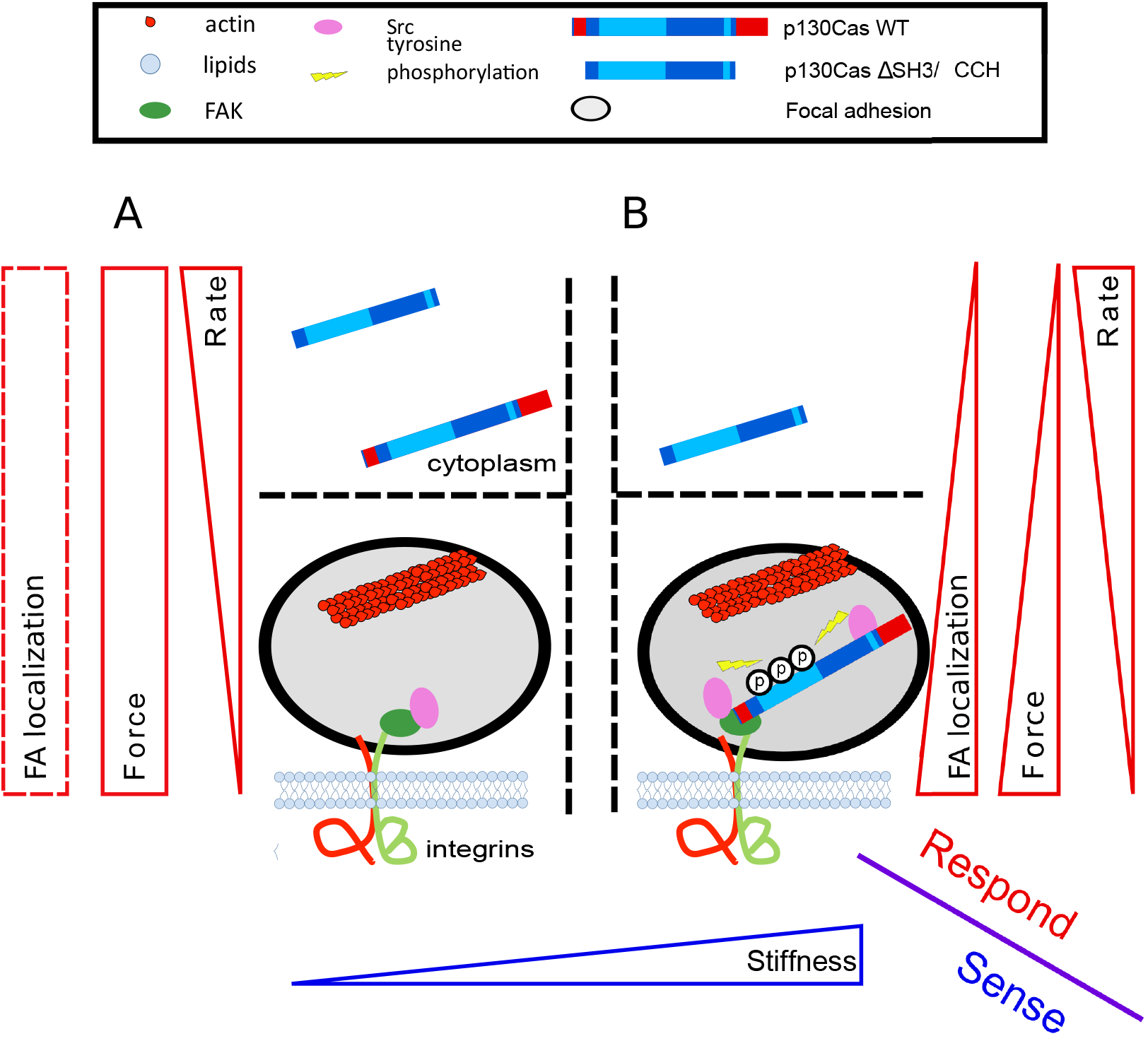
Mechanosensing and force exertion response at FAs via p130Cas. Cells sense (blue) and respond (red) to the stiffness in their environment via FAs in a p130Cas-dependent manner. In response to stiffness, p130Cas localizes and changes force magnitude and rate. On soft substrates (A), p130Cas remains in the cytoplasm, resulting in a loss of docking with multiple molecular players and an overall decrease in tyrosine phosphorylation in FAs. On stiffer substrates (B), p130Cas localizes well to FAs, increasing the probability of tyrosine phosphorylation within FAs, which leads to higher force magnitude, and lower force exertion rates.

## Conclusion

The data presented here provide evidence for the constitutive biochemical-mechanical link through which p130Cas modulates cellular force exertion depending on extracellular stiffness. These results might aid to the understanding of how p130Cas influences processes such as metastasis and invasiveness in cancer, where p130Cas has previously been noted (12; 15). Indeed, in MCF10A-CA1d breast carcinoma cells, it has been shown that p130Cas localizes to podosomes that form preferentially on stiffer substrates. In these latter experiments, the percentage of area actively degraded by the cell on hard substrates compared to soft substrates was directly influenced by p130Cas expression (39). We further speculate that our proposed molecular-mechanical model for p130Cas mechanosensing, as summarized in Fig.7, might be relevant not only in FAs, but also other ECM-cell contacts such as invadosomes. Further investigations into p130Cas’s influence on force exertion at invadosomes would, therefore, be of future interest.

## Acknowledgments

The authors gratefully acknowledge the help of Karin Jansen and GijsjeKoenderink at AMOLF for help with the PA gel stiffness measurements. We also thank Wim Pomp for help with the focus-hold and the automatedXY-positioning system. We thank the laboratory of Kees Storm at TU Eindhoven for help with finite element modelling of pillar deflections. We thank Steve Hanks at Vanderbilt University for Cas MEF cell lines. We also thank our other collaborators within the Mechanobiology NLconsortium for their insightful comments and suggestions. This work was supported by funds from the Netherlands Organization for Scientific Research (NWO-FOM) within the program on Mechanosensing and Mechanotransduction by Cells (FOM L1712M).

## Author contributions

Designed research: HvH,DMD,EHD,TS

Performed research: HvH,DMD,HEB

Contributed analysis tools / analyzed data: HvH,DMD,HEB,TS Wrote the paper: HvH,DMD,EHD,TS

## Supporting information

**Figure S1:**
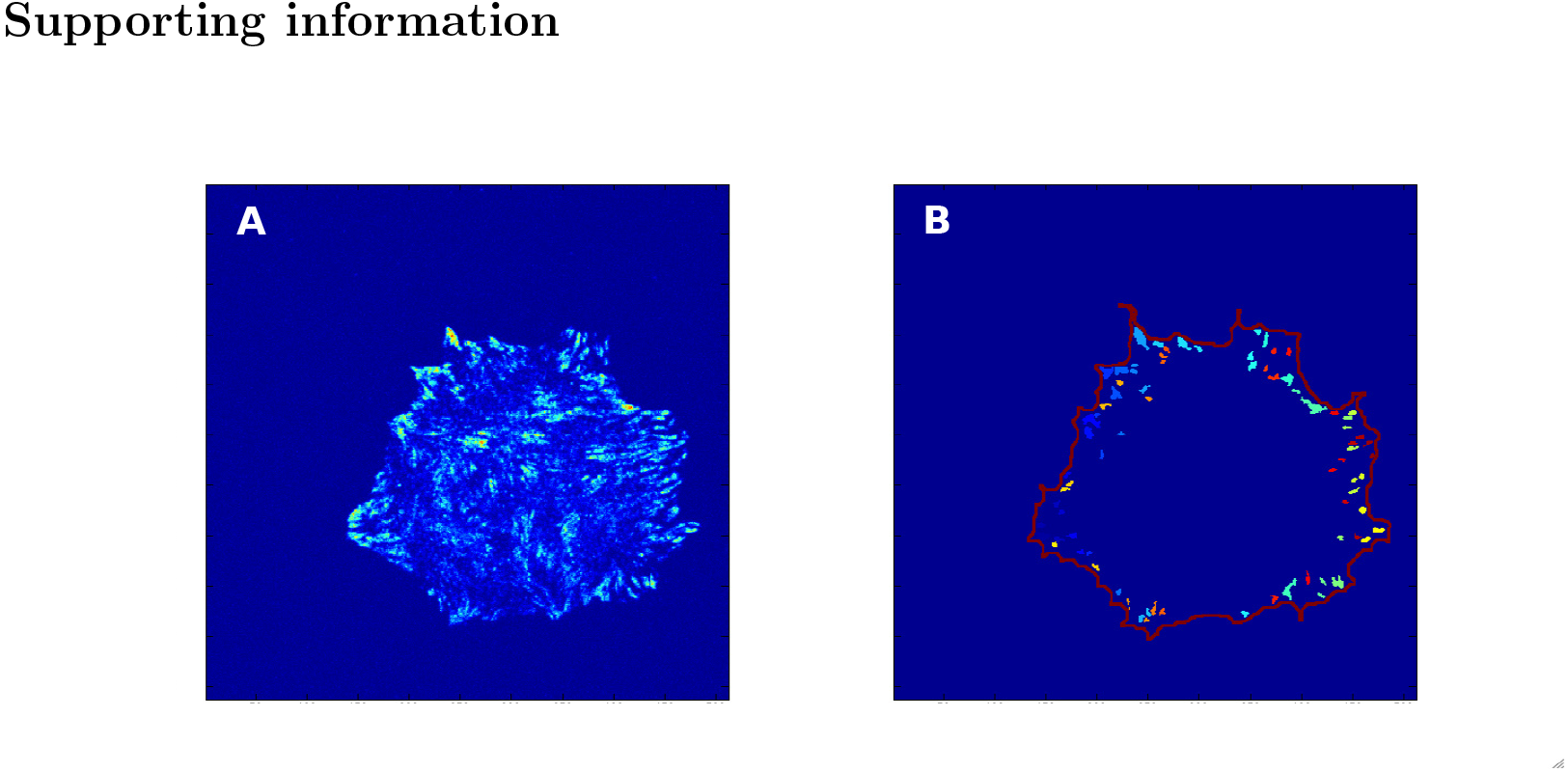
Matlab analysis for cell area and FA characterization. (A) Fluorescence image, and (B) FA-detection panel. The analysis yields information on cell-area, FA-area, FA-orientation from the binary image shown in (B). For clarity, the FAs are colour-coded.

**Figure S2:**
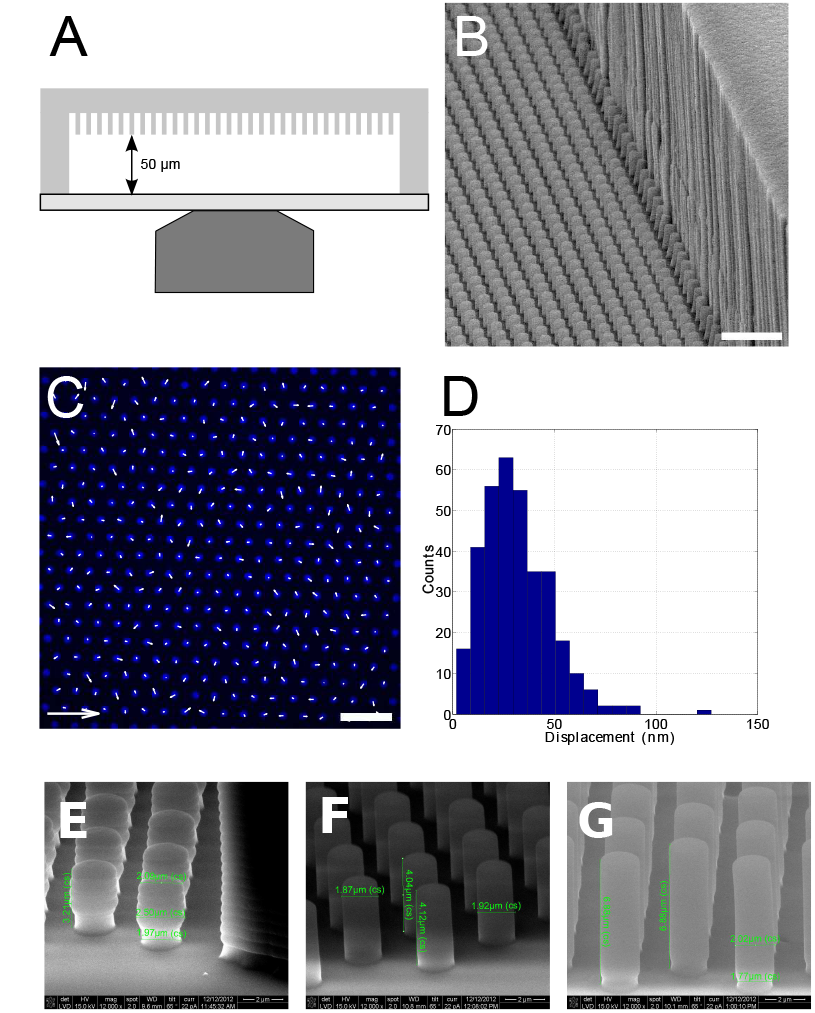
PDMS micropillar arrays for cellular force measurements. (A) Arrangment of the micropillar arrays on the optical microscope. (B) Electron-microscopy image showing the pillar array and the integrated spacer structure to the right. (C) Optical image of an undeflected micropillar array. The top of the pillars were stamped with fluorescence-labeled fibronection. The displacements shown as arrows reflect the positional accuracy to detect the center-of-mass of a single pillar. (D) Absolute pillar deflections of an undeflected array. Pillars were detected with a positional accuracy of 30 nm. (E) Micropillars with an effective stiffness of 12 kPa had a height of 6.9*μ*m and a diameter of 2.0*μ*m. (F) Those with an effective stiffness of 47 kPa had an average height of 4.1*μ*m and a post a diameter of 1.9*μ*m. (G) Micropillars at the 137 kPa stiffness had an average diameter of 2.1*μ*m and a height of 3.2*μ*m.

**Figure S3:**
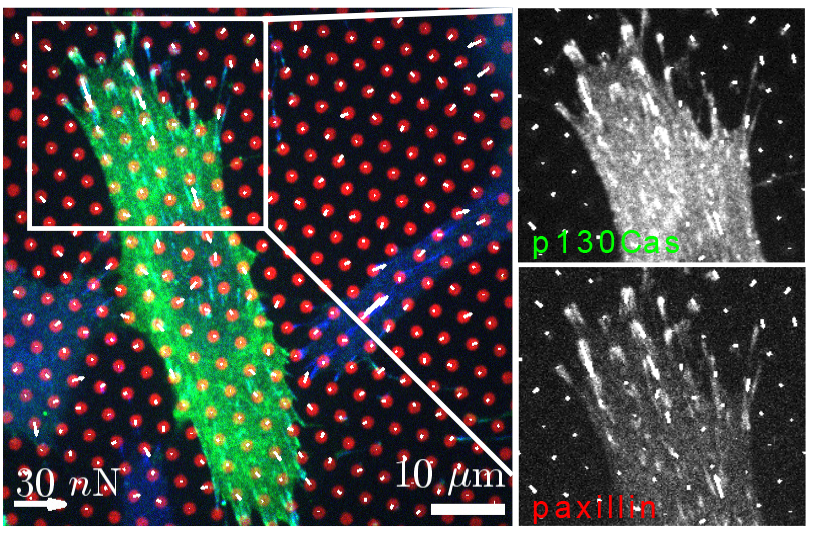
p130Cas localization to pillars on which force is exerted. 47 kPA micropillars were stamped with fibronectin-Alexa647 (red). p130Cas-YFP (green) co-localizes with paxillin-Alexa405 (blue) in FAs where force is exerted. Insets of the p130Cas (top) and the paxillin (bottom) channel in grayscale are at 125% of the original image magnification.

**Figure S4:**
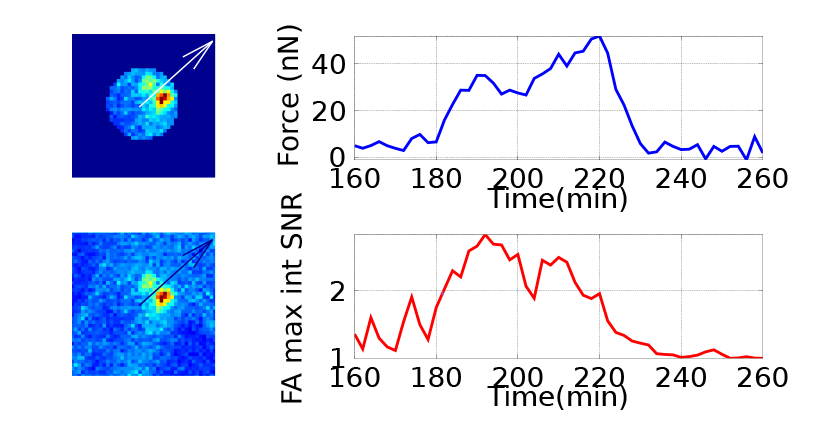
Quantification of p130Cas fluorescence on a single pillar to which a force is exerted. p130Cas fluorescence was integrated over a radius of 1.2 *μ*m around the pillar (masked image in upper left and unmasked image in lower left). The force over time (upper right) exerted on this pillar was correlated to the fluorescence increase i.e. p130Cas localization (lower right). It becomes evident that both signals display a strong time-correlated dynamics.

**Figure S5:**
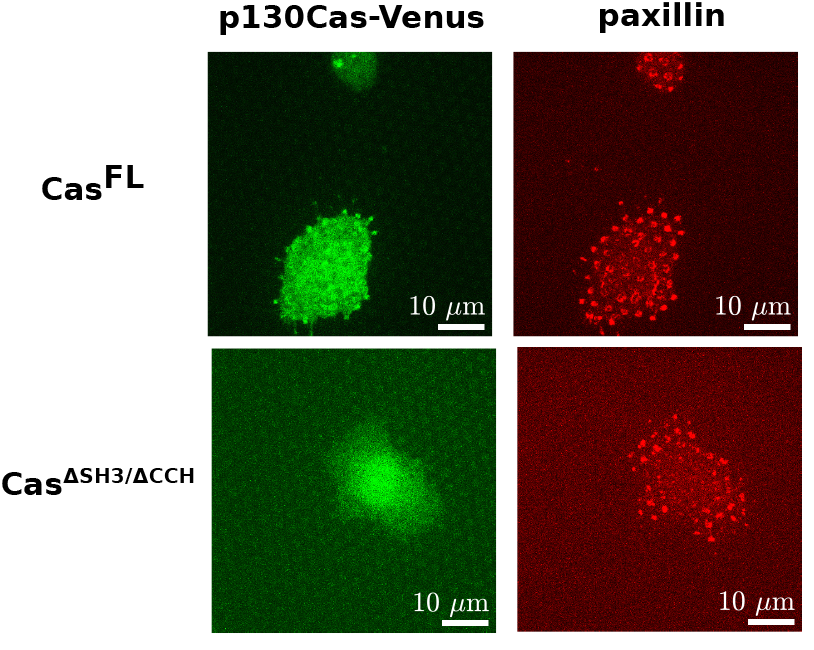
CAS^∆SH3/∆SSH^failed to localize to focal adhesions. On 137 kPa pillar arrays CAS^FL^(top, left) localized to focal adhesions as visualized by labeling paxillin (top, right). In comparison CAS^∆SH3/∆SSH^stayed cytosolic (bottom).

*Video S1.* **Movie 1** shows pillars with an effective stiffness of 12 kPa. Several cells migrate randomly in the field-of-view and exert transient forces of 1-20 nN. Only forces that were above the detection sensitivity of the system > 240 pN where taken into account for force analysis.

*Video S2.* **Movie 2** shows pillars with an effective stiffness of 47 kPa. Several cells migrate randomly in the field-of-view and exert transient forces of 5-30 nN. Only forces that were above the detection sensitivity of the system > 980 pN where taken into account for force analysis.

*Video S3.* **Movie 3** shows pillars with an effective stiffness of 137 kPa. Several cells migrate randomly in the field-of-view and exert transient forces of 10-60 nN. Only forces that were above the detection sensitivity of the system > 3.0 nN where taken into account for force analysis.

**Table S1.** Dependence of the stiffness on the ratio acrylamide to bis-acrylamide in. The shear moduli G was quantified in rheometer measurements. Assuming a Poisson ratio of *ν* = 0.5 the Young’s modulus E was calculated from the shear modulus by *E* = 2 (1 + *ν*) *G*. In this paper stiffness is consistently reported in Youngs’ modulus E.

| Acrylamide (%) | Bis-acrylamide (%) | $G$ (kPa) | $E$ (kPa) |
| --- | --- | --- | --- |
| 7.5 | 0.2 | 9.3 | 28 |
| 7.5 | 0.5 | 14 | 42 |
| 12 | 0.6 | 29 | 87 |
| 20 | 0.7 | 74 | 222 |

**Table S2.**
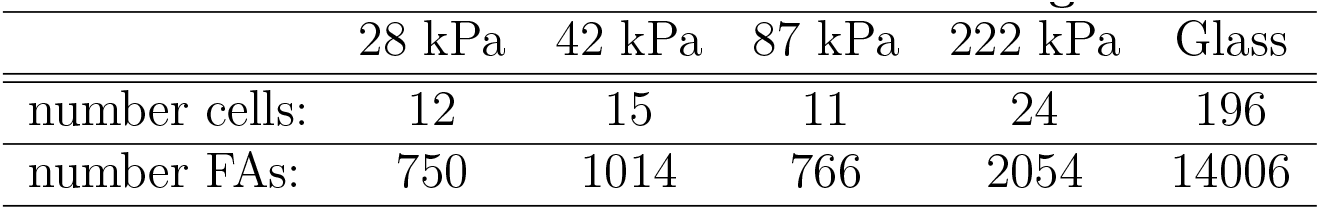
Number of measurements on PA gels.

**Table S3.** Number of measurements on micropillar arrays.

|  | 12 kPa | 47 kPa | 137 kPa |
| --- | --- | --- | --- |
| number force curves (Cells): | 153(12) | 243(20) | 119(13) |
| number fluorescence/force comparison (Cells): | 423(18) | 266(26) | 290(21) |
| number timelags (Cells): |  |  | 44(3) |

## References

[1] Moore, S. W., P. Roca-Cusachs, and M. P. Sheetz, 2010. Stretchy proteins on stretchy substrates: the important elements of integrin-mediated rigidity sensing. Developmental cell 19:194–206.

[2] Polacheck, W. J., R. Li, S. G. Uzel, and R. D. Kamm, 2013. Microfluidic platforms for mechanobiology. Lab on a Chip 13:2252–2267.

[3] Swift, J., I. L. Ivanovska, A. Buxboim, T. Harada, P. D. P. Dingal, J. Pinter, J. D. Pajerowski, K. R. Spinler, J.-W. Shin, M. Tewari, et al., 2013. Nuclear lamin-A scales with tissue stiffness and enhances matrix-directed differentiation. Science 341:1240104.

[4] Sochol, R. D., A. T. Higa, R. R. Janairo, S. Li, and L. Lin, 2011. Unidirectional mechanical cellular stimuli via micropost array gradients. Soft Matter 7:4606–4609.

[5] DuFort, C. C., M. J. Paszek, and V. M. Weaver, 2011. Balancing forces: architectural control of mechanotransduction. Nature reviews Molecular cell biology 12:308–319.

[6] Swartz, M. A., and A. W. Lund, 2012. Lymphatic and interstitial flow in the tumour microenvironment: linking mechanobiology with immunity. Nature Reviews Cancer 12:210–219.

[7] Dupont, S., L. Morsut, M. Aragona, E. Enzo, S. Giulitti, M. Cordenonsi, F. Zanconato, J. Le Digabel, M. Forcato, S. Bicciato, et al., 2011. Role of YAP/TAZ in mechanotransduction. Nature 474:179–183.

[8] del Rio, A., R. Perez-Jimenez, R. Liu, P. Roca-Cusachs, J. M. Fernandez, and M. P. Sheetz, 2009. Stretching single talin rod molecules activates vinculin binding. Science 323:638–641.

[9] Zaidel-Bar, R., and B. Geiger, 2010. The switchable integrin adhesome. Journal of cell science 123:1385–1388.

[10] Prager-Khoutorsky, M., A. Lichtenstein, R. Krishnan, K. Rajendran, A. Mayo, Z. Kam, B. Geiger, and A. D. Bershadsky, 2011. Fibroblast polarization is a matrix-rigidity-dependent process controlled by focal adhesion mechanosensing. Nature cell biology 13:1457–1465.

[11] Kim, D.-H., S. B. Khatau, Y. Feng, S. Walcott, S. X. Sun, G. D. Longmore, and D. Wirtz, 2012. Actin cap associated focal adhesions and their distinct role in cellular mechanosensing. Scientific reports 2.

[12] Tikhmyanova, N., J. L. Little, and E. A. Golemis, 2010. CAS proteins in normal and pathological cell growth control. Cellular and molecular life sciences 67:1025–1048.

[13] Van Der Flier, S., A. Brinkman, M. P. Look, E. M. Kok, M. E. Meijer-van Gelder, J. G. Klijn, L. C. Dorssers, and J. A. Foekens, 2000. Bcar1/p130Cas protein and primary breast cancer: prognosis and response to tamoxifen treatment. Journal of the National Cancer Institute 92:120–127.

[14] Huang, W., B. Deng, R.-W. Wang, Q.-Y. Tan, Y. He, Y.-G. Jiang, and J.-H. Zhou, 2012. BCAR1 protein plays important roles in carcinogenesis and predicts poor prognosis in non-small-cell lung cancer. PloS one 7:e36124.

[15] Barrett, A., C. Pellet-Many, I. C. Zachary, I. M. Evans, and P. Frankel, 2013. p130Cas: a key signalling node in health and disease. Cellular signalling 25:766–777.

[16] Meenderink, L. M., L. M. Ryzhova, D. M. Donato, D. F. Gochberg, I. Kaverina, and S. K. Hanks, 2010. P130Cas Src-binding and substrate domains have distinct roles in sustaining focal adhesion disassembly and promoting cell migration. PLoS One 5:e13412.

[17] Webb, D. J., K. Donais, L. A. Whitmore, S. M. Thomas, C. E. Turner, J. T. Parsons, and A. F. Horwitz, 2004. FAK–Src signalling through paxillin, ERK and MLCK regulates adhesion disassembly. Nature cell biology 6:154–161.

[18] Donato, D. M., L. M. Ryzhova, L. M. Meenderink, I. Kaverina, and S. K. Hanks, 2010. Dynamics and mechanism of p130Cas localization to focal adhesions. Journal of Biological Chemistry 285:20769–20779.

[19] Shin, N.-Y., R. S. Dise, J. Schneider-Mergener, M. D. Ritchie, D. M. Kilkenny, and S. K. Hanks, 2004. Subsets of the major tyrosine phosphorylation sites in Crk-associated substrate (CAS) are sufficient to promote cell migration. Journal of Biological Chemistry 279:38331–38337.

[20] Sawada, Y., M. Tamada, B. J. Dubin-Thaler, O. Cherniavskaya, R. Sakai, S. Tanaka, and M. P. Sheetz, 2006. Force sensing by mechanical extension of the Src family kinase substrate p130Cas. Cell 127:1015–1026.

[21] Lu, C., F. Wu, W. Qiu, and R. Liu, 2013. P130Cas substrate domain is intrinsically disordered as characterized by single-molecule force measurements. Biophysical chemistry 180:37–43.

[22] Hotta, K., S. Ranganathan, R. Liu, F. Wu, H. Machiyama, R. Gao, H. Hirata, N. Soni, T. Ohe, C. W. Hogue, et al., 2014. Biophysical properties of intrinsically disordered p130Cas substrate domainimplication in mechanosensing. PLoS computational biology 10:e1003532.

[23] Wang, X., and T. Ha, 2013. Defining single molecular forces required to activate integrin and notch signaling. Science 340:991–994.

[24] Yeung, T., P. C. Georges, L. A. Flanagan, B. Marg, M. Ortiz, M. Funaki, N. Zahir, W. Ming, V. Weaver, and P. A. Janmey, 2005. Effects of substrate stiffness on cell morphology, cytoskeletal structure, and adhesion. Cell motility and the cytoskeleton 60:24–34.

[25] Balcioglu, H. E., H. van Hoorn, D. M. Donato, T. Schmidt, and E. H. Danen, 2015. Integrin expression profile modulates orientation and dynamics of force transmission at cell matrix adhesions. Journal of Cell Science jcs–156950.

[26] van Hoorn, H., R. Harkes, E. M. Spiesz, C. Storm, D. van Noort, B. Ladoux, and T. Schmidt, 2014. The nanoscale architecture of force-bearing focal adhesions. Nano letters 14:4257–4262.

[27] Ghibaudo, M., A. Saez, L. Trichet, A. Xayaphoummine, J. Browaeys, P. Silberzan, A. Buguin, and B. Ladoux, 2008. Traction forces and rigidity sensing regulate cell functions. Soft Matter 4:1836–1843.

[28] Trichet, L., J. Le Digabel, R. J. Hawkins, S. R. K. Vedula, M. Gupta, C. Ribrault, P. Hersen, R. Voituriez, and B. Ladoux, 2012. Evidence of a large-scale mechanosensing mechanism for cellular adaptation to substrate stiffness. Proceedings of the National Academy of Sciences 109:6933–6938.

[29] Honda, H., H. Oda, T. Nakamoto, Z.-i. Honda, R. Sakai, T. Suzuki, T. Saito, K. Nakamura, K. Nakao, T. Ishikawa, et al., 1998. Cardiovascular anomaly, impaired actin bundling and resistance to Src-induced transformation in mice lacking p130Cas. Nature genetics 19:361–365.

[30] Fu, J., Y.-K. Wang, M. T. Yang, R. A. Desai, X. Yu, Z. Liu, and C. S. Chen, 2010. Mechanical regulation of cell function with geometrically modulated elastomeric substrates. Nature methods 7:733–736.

[31] Han, S. J., K. S. Bielawski, L. H. Ting, M. L. Rodriguez, and N. J. Sniadecki, 2012. Decoupling substrate stiffness, spread area, and micropost density: a close spatial relationship between traction forces and focal adhesions. Biophysical journal 103:640–648.

[32] Mih, J. D., A. Marinkovic, F. Liu, A. S. Sharif, and D. J. Tschumperlin, 2012. Matrix stiffness reverses the effect of actomyosin tension on cell proliferation. Journal of cell science 125:5974–5983.

[33] Balaban, N. Q., U. S. Schwarz, D. Riveline, P. Goichberg, G. Tzur, I. Sabanay, D. Mahalu, S. Safran, A. Bershadsky, L. Addadi, et al., 2001. Force and focal adhesion assembly: a close relationship studied using elastic micropatterned substrates. Nature cell biology 3:466–472.

[34] Bae, Y. H., K. L. Mui, B. Y. Hsu, S.-L. Liu, A. Cretu, Z. Razinia, T. Xu, E. Pure, and R. K. Assoian, 2014. an FAK-Cas-Rac-lamellipodin signaling module transduces extracellular matrix stiffness into mechanosensitive cell cycling. Science signaling 7:ra57–ra57.

[35] Machiyama, H., H. Hirata, X. K. Loh, M. M. Kanchi, H. Fujita, S. H. Tan, K. Kawauchi, and Y. Sawada, 2014. Displacement of p130Cas from focal adhesions links actomyosin contraction to cell migration. Journal of cell science 127:3440–3450.

[36] Ballestrem, C., N. Erez, J. Kirchner, Z. Kam, A. Bershadsky, and B. Geiger, 2006. Molecular mapping of tyrosine-phosphorylated proteins in focal adhesions using fluorescence resonance energy transfer. Journal of cell science 119:866–875.

[37] Janostiak, R., J. Brabek, V. Auernheimer, Z. Tatarova, L. A. Lautscham, T. Dey, J. Gemperle, R. Merkel, W. H. Goldmann, B. Fabry, et al., 2014. CAS directly interacts with vinculin to control mechanosensing and focal adhesion dynamics. Cellular and molecular life sciences 71:727–744.

[38] Cho, S. Y., and R. L. Klemke, 2000. Extracellular-regulated kinase activation and CAS/Crk coupling regulate cell migration and suppress apoptosis during invasion of the extracellular matrix. The Journal of cell biology 149:223–236.

[39] Alexander, N. R., K. M. Branch, A. Parekh, E. S. Clark, I. C. Iwueke, S. A. Guelcher, and A. M. Weaver, 2008. Extracellular matrix rigidity promotes invadopodia activity. Current Biology 18:1295–1299.

